# Insecticide-free trapping bed-net can mitigate insecticide resistance threat in malaria vector control strategies

**DOI:** 10.1101/2020.02.06.937623

**Authors:** Chouaibou S. Mouhamadou, Kun Luan, Behi Kouadio Fodjo, Andre West, Marian McCord, Charles S. Apperson, R. Michael Roe

## Abstract

Mosquito-borne malaria kills 429,000 people each year with the problem acute in sub-Saharan Africa. The successes gained with long-lasting pyrethroid treated bed-nets is now in jeopardy because of wide-spread, pyrethroid-resistance in mosquitoes. Using crowd modeling theory normalized for standard bed-net architecture, we were able to design an attract-trap-kill mechanism for mosquitoes that does not require insecticides. Using three-dimensional polyester knitting and heat fixation, trap funnels were developed with high capture efficacy, no egression under worst-case laboratory conditions, and greater durability than current bed-nets sold. Field testing in Africa in WHO huts with Gen1-3 T (trap)-Nets validated our model, and as predicted, Gen3 had the highest efficacy with a 4.3-fold greater trap-kill rate with no deterrence or repellency compared to Permanet 2.0, the most common bed-net in Africa. A T-Net population model was developed based on field data to predict community level mosquito control compared to a pyrethroid bed-net. This model showed the Gen3 T-Net under field conditions in Africa against pyrethroid resistant mosquitoes was 12.7-fold more efficacious than single chemical, pyrethroid treated nets, demonstrating significantly greater mosquito control using bed-nets without insecticides.

## Introduction

Malaria is the leading cause of morbidity and mortality in Sub-Saharan Africa with 212 million cases estimated by the WHO and about 429,000 deaths each year^1^. Malaria prevention is mainly based on vector (mosquito) control, using insecticide indoor residual spraying (IRS) or in long-lasting insecticide-impregnated mosquito bed-nets (LLINs)^2^. The efficacy of these measures depends primarily on the susceptibility of the mosquito to insecticides. An estimated 663 million cases of malaria have been averted in sub-Saharan Africa since 2001 as a result of the scale-up of malaria control interventions; 69% of these cases were a direct result of the use of LLINs^3^. Unfortunately, mosquitoes have become resistant to insecticides in recent years^4–6^ (including pyrethroids, the only class of insecticides presently used in bed-nets), threatening the efficacy of insecticide-based vector control programs^7–9^. This situation is problematic and highlights the urgent need to develop new strategies to mitigate development and spread of insecticide resistance and preserve the efficacy of currently available insecticides and malaria control interventions.

Development of a new, bed-net insecticide chemistry that is safe to human exposure every sleep cycle, that must survive for years of human handling and washing, and will not promote mosquito resistance and cross-resistance is a major challenge^10^. Alternative solutions include the reformulation of agricultural chemicals for vector control with completely different modes of action from the common public health insecticides. However, it is difficult with these solutions to account for prior and future exposure of mosquito larvae to pesticides used in farming resulting again in resistance^11^. All of these challenges have led us to focus on a different approach, a mechanical killing solution for mosquito control.

It is well known that mosquitoes are attracted to carbon dioxide and other human odors, and these odorants emitted from the body are warmer than ambient air and rise from its source^12^. Infrared video tracking has now shown that about 75% of mosquitoes follow this odorant plume and are attracted to the top surface of a bed-net in use^13,14^. Hence, we hypothesized that mosquito movement in the absence of visual and odorant cues would be random. If we then treat mosquitoes as ideal particles in an ideal gas, we can use the Maxwell-Boltzmann distribution^15^ to develop a mathematic model for the construction of an insecticide-free trapping bed-net (T-Net) to capture and hold host-seeking mosquitoes during and after a sleep cycle. The T-Net is a bed-net with two compartments, including (i) a lower sleeping compartment and (ii) an upper mosquito trap containing cone-like funnels as mosquito entry points into the trap space. 3-D knitting was used for cone construction. Proof of concept was investigated in the laboratory and then under field conditions with wild-type, resistant *An. gambiae* mosquitoes in Tiassale, Côte d’Ivoire (Africa) to verify our model. From the field data, we developed a second model to describe community level mosquito control by the T-Net versus the most common bed-net deployed in Africa.

## Results

### Crowd model applied to mosquito trapping

In our effort to develop a non-insecticidal bed net that kills mosquitoes by trapping, studies were conducted to model mosquito interactions with a bed net designed for trapping. The model if predictive of field conditions could be used to evaluate different trap designs and make a better mosquito trap. The model could also provide a better understanding of how mosquitoes interact with bed-nets. If we treat a mosquito as an ideal gas particle, and the flight track is assumed to be random, the Maxwell-Boltzmann distribution^15^ can be used for defining the random flight of mosquitoes at different velocities. This can further be applied to the size dimensions in the context of a standardized WHO approved, experimental research hut^16^ as shown in Tiassale, Cote d’Ivoire (Africa) where our field work was conducted (Fig. 1a-b). Fig. 1b diagrams the process by which mosquitoes can fly from the interior of the hut through knitted cones into a trap compartment mounted above the sleeping compartment. Mosquitoes that are in the trap compartment are blocked from the sleeper by the trap bottom, which is also the sleep compartment roof. Once the mosquitoes are trapped, they become exhausted and quiescent, and die from dehydration a few hours later.

**Figure 1.**
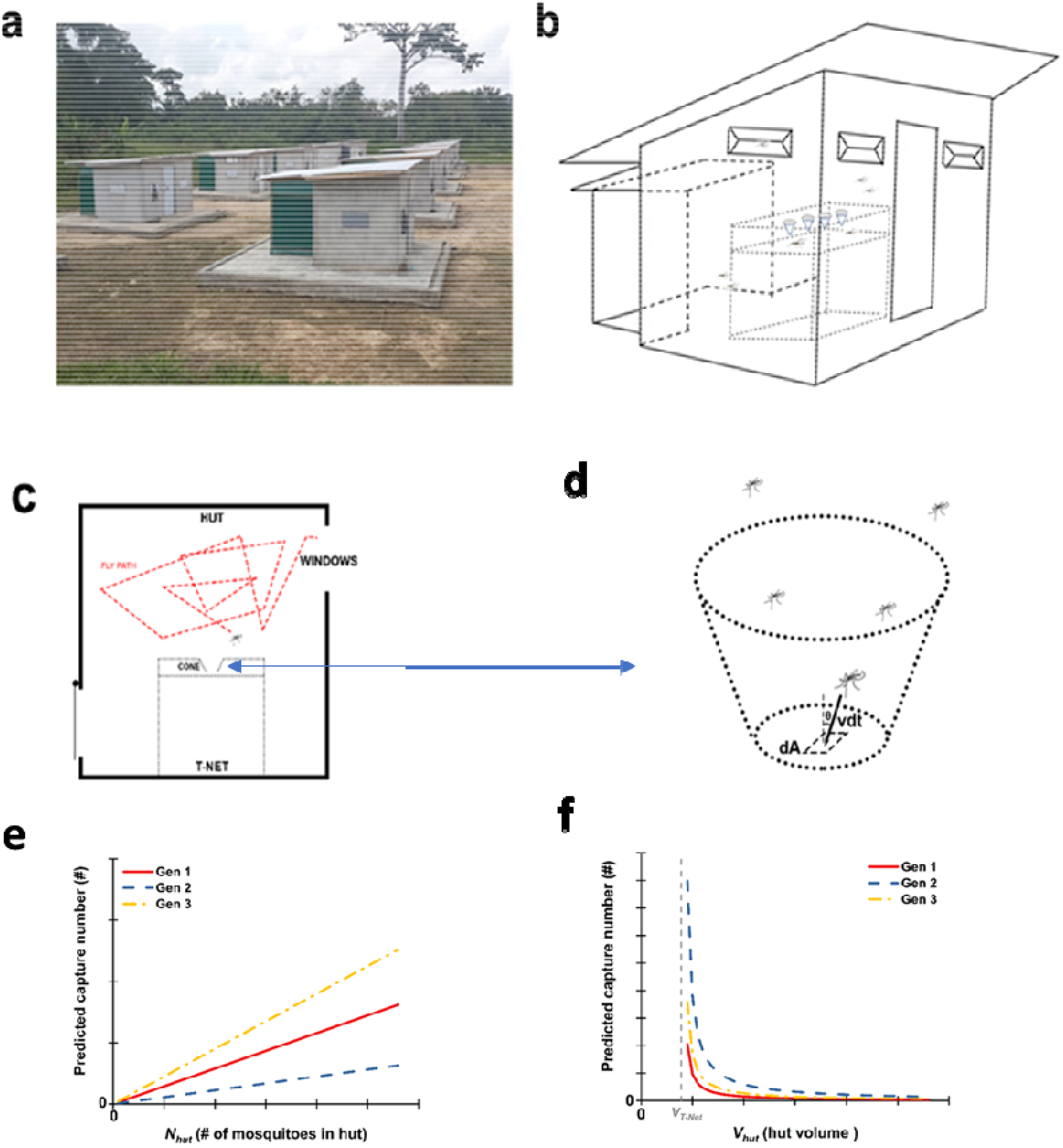
Principle and model of the T-Net: (a): picture of experimental huts in the region of Tiassale in Cote d’Ivoire (Africa), where the field trials were conducted; (b): schematic of hut. (c): schematic of mosquito flight track in the hut; (c): unit area on the bottom of T-Net cone that captures mosquitoes, (e): predicted capture number for Gen 1, Gen 2 and Gen 3 T-Nets (see Fig. 2) with an uncertain number of mosquito flying into the hut, (f) the predicted capture number for Gen 1, Gen 2 and Gen 3 T-Nets when hut volume increases and the number of mosquitoes flying into the hut is constant.

Knitted cones were developed as access points to the trap compartment. They can easily be constructed in any shape and size for prototyping, are easily sewn into bed-nets, and can fold into a flat, compact surface critical for storing and shipping of bed-nets. Cones (Fig. 2) were knitted on a flat, computerized knitting machine, SWGN2 (Shima Seiki, Japan), where their dimensions for prototyping are easily changed by standard program options. This gauge of machine was selected so that the resulting cone fabric would have openings between loops similar to those of traditional bed-nets. Similar to the construction of stockings or socks, cones were knitted in-the-round to make a 3D shape (Fig. 2). Use of this construction method allowed us to alter the shape, depth, width, and height in all sections of the cone to meet our design demands. The knitted cones had three main parts: a large opening at the mouth (which was designed to allow for passage of mosquitoes into the cone), tapered cone walls (which were designed to direct the movement of mosquitoes into the trapping compartment), and the neck of the cone opening into the trapping compartment (the size of the neck was optimized to allow passage of mosquitoes and minimize egression from the trap). To prevent collapsing and flattening of the knitted cones, they were constructed with thermal bond fibers. This yarn is a polyester binder spun yarn in which each fiber is composed of a sheath part (co-polyester) and a core (regular polyester). The bursting strength and durability of these cones was compared to conventional bed-net material (see the Extended data) to make sure they would stand up to years of use.

**Figure 2:**
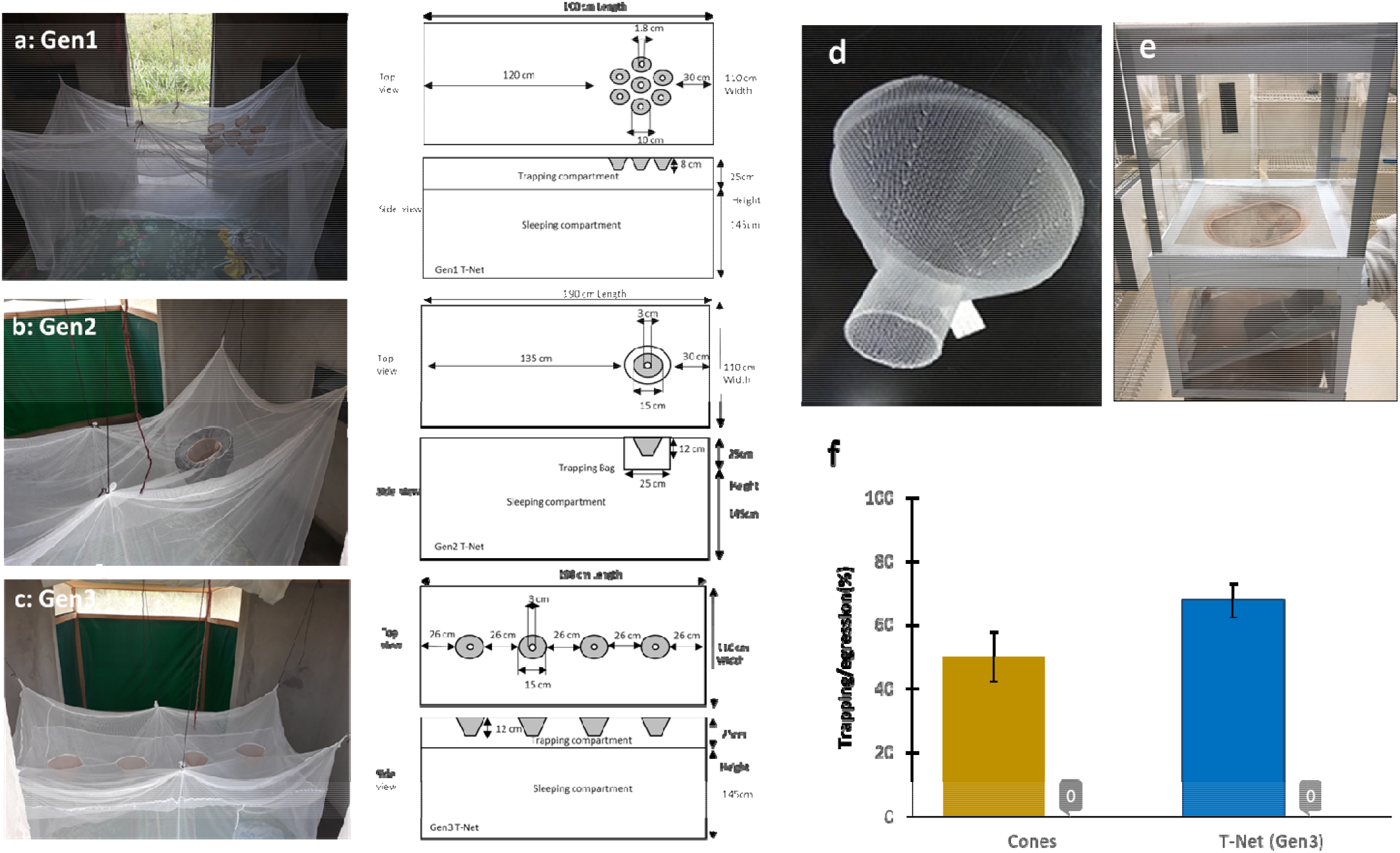
Different designs of T-Nets and proof of concept findings. (a): Gen1 T-Net made of a circular aggregate of seven small cones; (b): Gen2 T-Net made with one large cone fitted on a trapping bag; (c): Gen3 T-Net fitted with four large cones; (d): example of a knitted cone; (e): experimental tunnel apparatus to assess the trapping efficacy of knitted cones; (f) results obtained on tunnel test for the trapping efficiency of large cones (on the left) and for the Gen3 T-Net in a laboratory hut experiment (on the right). The histograms represent the trapping rates, the bars on top are the confidence intervals, and “0” are the exit rates (=egression) in absence of host cues.

In respect to model development, we assumed that (i) the flight track of mosquitoes is random (Fig. 1c); (ii) the diameters of the cone into the trap compartment are large enough not to affect the flight path of other mosquitoes; and (iii) the trapping does not perturb the flight velocity of the remaining mosquitoes in the hut. Fig. 1d assumes a hole area *dA*, a mosquito that is at a distance *vdt* d from the hole, and a mosquito moving at a speed *v* at an angle *θ* from the normal surface toward the area *dA*. All mosquitoes within a parallelepiped volume around the *dA* area moving toward the hole with speed *v* will pass through the top opening of the cone in the time interval *dt*. Therefore, the total number of mosquitoes flying through the *dA* area in the time interval *dt* is:

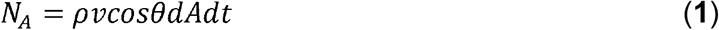

where *ρ* is the mosquito density in the hut.

Assuming the distribution of individual mosquito velocities obeys the Maxwell-Boltzmann distribution, which was first defined and used for describing particle speeds in idealized gases, one integrated expression of the distribution relates particle density with average velocity. Similarly, the average trapped mosquito number per area per time is then:

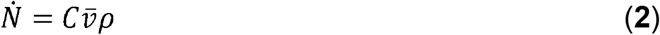

where 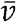 is the average flight speed, *ρ* is the mosquito density in the hut and *C* is a constant determined by the active behavior of the mosquito (1/4 in ideal gas theory).

As mentioned above, most of mosquitoes fly to the top of the bed-net and thus the work volume is:

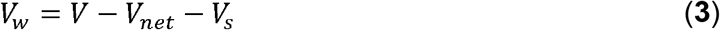

where *V* is the hut volume, *V_net_* is the bed-net volume and *V_s_* is the volume surrounding the bed-net except the volume on the top of the bed-net.

Assuming mosquitoes are distributed evenly on the top of the bed-net, the trapped mosquito number for the T-Net in the testing time *t* is:

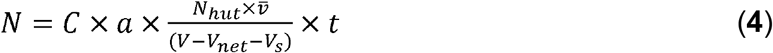

where *N_hut_* is the total mosquito number in the hut, *C* is a constant and *a* is the bottom area of the cone.

Thus, the T-Net model relates trapping number with container volume and flight velocity. If the number of mosquitoes in a hut is a variable, and other parameters are constant, the predicted results of equation (**4**) are shown in Fig. 1e; if *V_hut_* is a variable, the predicted trapping number for the T-Net is shown in Fig. 1e for three different versions of the net that we investigated for model development and field validations (Gen1-3 prototypes shown in Fig. 2).

### T-Net construction and Model Prediction

The attraction of mosquitoes through the funnels into the trap compartment is hypothesized to be from insect attraction to carbon dioxide and other odors from human respiration. If this is the case, funnels should only be needed on the head end of the net. Based on this assumption, we designed and constructed the Gen1 T-Net with a circular aggregate of 7 cones (each cone having a 10 cm large opening, an 8 cm depth and 1.8 cm small diameter); these cones were sewn around the midline of the long axis of the trap roof (a 25 cm deep trap compartment) 30 cm from the sleeping end of the net (shown in Fig. 2a). The model (**4**) predicted trapping rates shown by the regression curves in Fig. 1e,f. To potentially increase the trapping rate and add additional functionality, a Gen2 T-Net was constructed (Fig. 2b). In this case, one large cone 15 cm in the large diameter, 12 cm deep and 3 cm in the small diameter, was fitted into a cylindrical trapping bag 25 cm deep and 25 cm in diameter fixed as before on the bed-net roof 30 cm from the edge. The idea of using a trapping bag and a single cone was to reduce material costs for construction, make it possible to reduce the head space between the sleeper and the bed net roof (effectively increasing the odorant concentration on the bed net top) and adding practical functionality to the bed net design. The Gen2 net design could be used to retrofit any non-trap bed net including those already deployed in homes. From equation (**4**), we predicted that this actually reduced the catch rate as compared to Gen1 (Fig. 1e,f).

Intuitively, having the cones just above the head of the sleeper should produce the maximum trap rate. At the same time, reducing the overall cross-sectional area of the small openings of the cones and/or the number of cones might reduce the catch rate but also reduce the egression rate once mosquitoes were captured. There is also the possibility that odorants from the subject’s body could be in play in trapping and spreading cones in different areas on the net top, might improve capture rates. To address the above in part, we increased the number of cones from 1 in Gen2 (Fig. 2b) to 4 in Gen3 (Fig. 2c). The Gen3 T-Net had four large cones (15 cm large diameter, 12 cm deep and 3 cm small diameter; same dimensions as Gen2), where the cones were aligned lengthwise along the centerline of the bed-net with the cones spaced 26 cm apart. All cones opened on a trapping box of 25 cm deep (Fig. 2c). Interesting, our model predicted an increase in the trapping rate over that for Gen1 and Gen 2 (Fig. 1e,f).

Our model (**4**) predicts capture rates but not the possibility of egression once mosquitoes are trapped. Therefore, laboratory bioassays were conducted with lab reared Kisumu strain, *An. gamibae* to examine egression rates across the large cones in Gen3 where the model predicted the highest capture rate. A high capture rate would not be optimal if the large cone and especially 4 large cones in Gen3 favored a higher egression versus trap rate. Studies were conducted using a modified WHO tunnel tests^17^ as shown in Fig. 2e. The capture rate (with mosquitoes released into the top chamber and the subject arm as shown in the bottom chamber) was 49.5% after 2 hours. More important, however, the egression rate once the mosquitoes were captured was 0% 2 h after the arm was removed (Fig. 2f). These results suggested that even though we added a greater overall area for mosquito entry into the trap in Gen3, the increased capture rate predicted by our model will not be affected by increased regression. The tunnel tests were also a worst-case scenario where the host cues were removed, and the trap volume was much smaller than that for Gen3 trap compartment.

Before going to the field in Africa to validate our trapping net model, a laboratory test was conducted with the Gen3 T-Net (Fig. 2c) in a walk in incubator (same dimensions as field huts, see details in material and method section) and with a human subject under the net. Susceptible, laboratory reared *An. gambiae* Kisumu were released into the room after the subject entered the sleeping compartment of the T-Net. A total of 300, unfed adult selected host-seeking female mosquitoes were used in six replicates with an average of 50 mosquitoes per replicate (range = 41-85 females). The trapping rate in these studies ranged from 45.28% to 86.90% after 3 hours with an average capture rate of 67.7% (± 20.5%, 95% CI)(Fig. 2f). No mosquito egression from the trapping compartment was observed at 30 minutes, 1h, 2h, 6h and overnight) for any of the replicates (Fig. 2f). These results demonstrated proof of concept that the predicted best performing T-Net, Gen3, under laboratory conditions simulating WHO experimental hut conditions at our field site in Africa, were efficacious in mosquito capture and retention.

### Field performance of Gen1, Gen2 and Gen3 T-Nets against insecticide resistant, wild, free-flying mosquitoes

To fully validate our model (**4**) for predicting the trapping efficacy for the T-Net, field trials were conducted comparing Gen1, Gen2 and Gen3 bed-nets with a positive control, Permanent 2.0 (PN2.0)-LLIN (the most used LLIN in Africa). The other objective of the field trials was to evaluate the efficacy of the T-Nets compared to a popular LLIN (Permanet 2.0) and determine if the technology is reasonably practical as a mosquito control strategy or whether more research is needed to further optimize the T-Net. These studies were conducted in WHO-approved experimental huts^16^ in the locality of Tiassale where wild *An. gambiae* malaria vectors are resistant to insecticides^18,19^ with human subjects sleeping under the nets each night. Three entomological parameters were measured: (i) the mean entry rate of mosquitoes per hut per night, (ii) the mean exit rate per hut each night, and (iii) the mean mortality rate per hut per night. For the T-Nets, we scored the trapped mosquitoes as alive or dead each morning but in practice, trapped mosquitoes are ecologically dead, i.e., they are quiescent, no longer able to blood feed and die typically before the next sleep cycle.

#### Mean entry number

A total of 53, 139, 175 and 156 *An. gambiae sl* mosquitoes were collected over 14 nights (representing 14 observations = 14 replicates), respectively, for PN2.0, Gen1, Gen2, Gen3 and (Table 1). The mean number of mosquitoes entering each hut per night was 3.8 (σ= 2.96) for PN 2.0 (the lowest entry rate in the field trial), followed by 9.9 (σ= 10.1) for Gen1, 12.5 (σ= 10.5) for Gen2 and 11.1 (σ= 8.3) for Gen3. The differences were not statistically significant between Gen1, Gen2 and Gen3, but a significant difference was observed between PN 2.0 and each of the T-Nets (P<0.05) (Fig. 3a).

**Table 1:** Results from field evaluations of Gen1, Gen2 and Gen3 T-Nets. Total caught, mean entry, mean exit and mean mortality rates per hut per night of *An. gambiae* mosquitoes after 14 observations during which the insecticide-free T-Nets were compared to Permanet 2.0 (PN2.0)-LLIN in Tiassale (Cote d’Ivoire) where mosquitoes are resistant to insecticides.

| Variable | # obs. | Total caught | Entry # |  | Exit rate (%) |  | Killing rate (%) |  |
| --- | --- | --- | --- | --- | --- | --- | --- | --- |
|  |  |  | Mean* | Std dev. | Mean* | Std dev. | Mean* | Std dev. |
| PN2.0 | 14 | 53 | 3.8a | 3.0 | 77.2a | 29.4 | 14.4a | 18.0 |
| Gen1 | 14 | 139 | 9.9b | 10.2 | 47.3b | 27.8 | 39.1b | 30.6 |
| Gen2 | 14 | 175 | 12.5b | 10.6 | 45.7b | 30.6 | 39.9b | 26.1 |
| Gen3 | 14 | 156 | 11.1b | 8.3 | 26.8b | 20.7 | 61.9c | 22.1 |

\*Values not sharing the same letters are statistically significant
#obs.: number of observations

**Figure 3:**
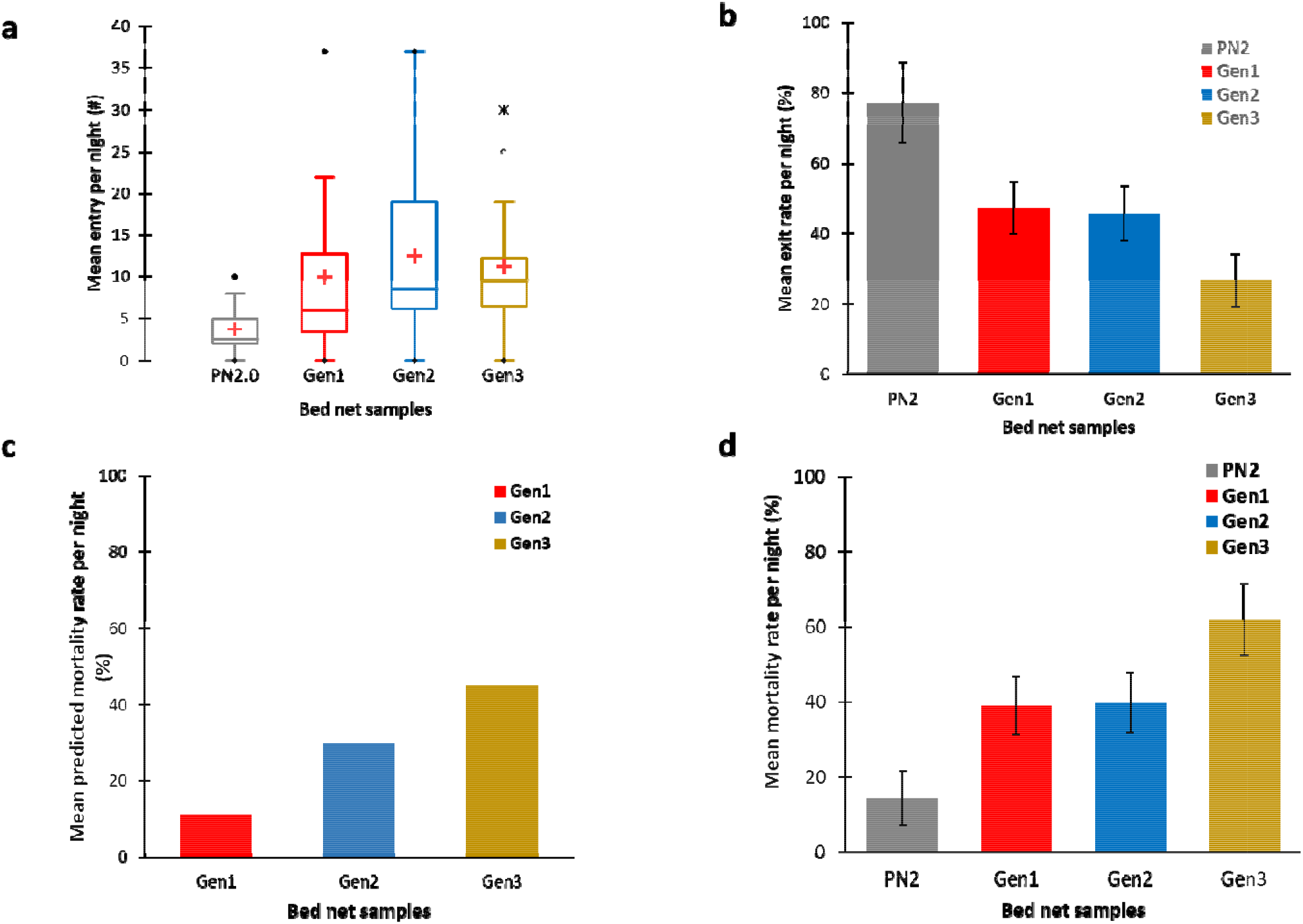
Performance of Gen1, Gen2 and Gen3 insecticide-free T-Nets in comparison to the Permanet 2.0 LLIN (PN2.0) in Tiassale (Cote d’Ivoire) where mosquitoes are resistant to insecticides. The nets were randomly allocated to four huts. Nets and sleepers were rotated each night. Up to 14 observations were made. (a) shows the mean number of mosquitoes entering each hut per night. (b) is the mean exit rate per hut per night, (c) is the killing rate of the T-Net; (d) is the mean mortality rate per hut per night (trapped = dead and ecologically dead). The bars on top of the histograms represent the confidence intervals.

#### Mean exit rate

As expected, the PN2.0 not only showed the lowest entry rate but also the highest mean exit rate of mosquitoes per night, 77.2% (σ = 29.4) and was statistically significantly different from that of Gen1, Gen2 and Gen3. In Gen1 and Gen2, we recorded a mean mosquito exit rate per night of 47.3% (σ= 27.7) and 45.7% (σ= 30.6), respectively, and in Gen3, the exit rate was 26.7% (σ= 20.7) which was the lowest exit rate. No statistically significant differences in exit rates was observed between the three T-Nets (Table 1; Fig. 3b).

#### Mean mortality rate

The lowest mortality rate (Fig. 3c) was occurred with PN2.0, 14.3% (σ= 17.9). It was statistically significantly lower than for all of the T-nets. The mortality rates for Gen1 and Gen 2 were 39.1% (σ = 30.5) and 39.8% (σ = 26.0), respectively, with no statistical significant differences between these two nets (P> 0.05). The highest mean mosquito mortality rate per hut per night was for Gen3 at 61.9.5% (σ= 22.1). This difference was higher than that of Gen1, Gen2 and PN.20 (P <0.05).

### T-Net model validation

Base on the tunnel test in Figure 2e, *C* in equation (**4**) was calculated by the condition parameters where the tunnel test cage dimension was 30cm in length, 30cm in width and 30cm in height. The cone trap diameter was 3cm.Thus, *C* is 0.00783. With these data, we predicted the trap number for Gen 1, Gen 2 and Gen 3, respectively, at 2.87, 1.41 and 5.02 corresponding to trapping rates of 28.9%, 11.3% and 45.06% (Fig. 4d). The relative trap efficiency between Gen1, Gen 2 and Gen3 calculated from the model was in agreement but underestimated the actual trap efficacy in the field trials. This was expected since the model did not take into consideration the attractants from the sleeper. For the Gen2, there was a much greater trap rate than predicted, suggesting that cone positioning above the sleeper head likely is responsible for a greater number of trapped mosquitoes than the other cones in Gen3. At the same time, the higher efficiacy of Gen3 where cones run along the long top axis of the net, confirms that odorants from other parts of the body increase trap efficacy.

**Figure 4:**
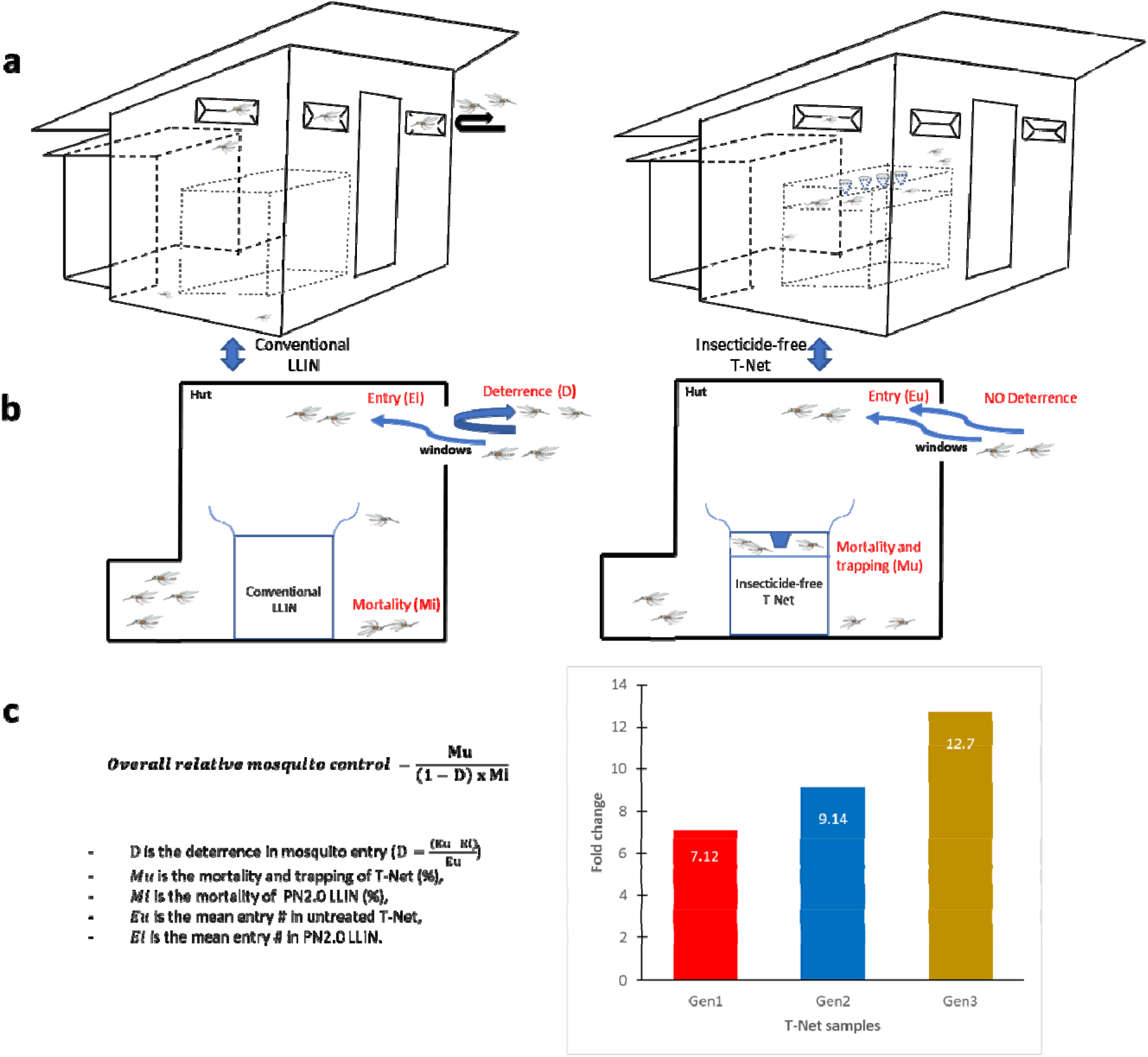
Overall relative mosquito control. (a): schematic illustration of the experimental hut with either a LLIN of a free-insecticide T-net; (b) highlights the different parameter used to define the model; (c) the overall relative mosquito control. The histograms showed the increased mortality of mosquitoes over PN2.0-LLIN.

### Model for community level, mosquito control using the T-Net

Gen1 and Gen2 T-Nets had a 2.7-fold greater kill rate than PN2.0 while Gen3 had a 4.3-fold greater kill rate than the permethrin treated (PN2.0) positive control. However, there is more to consider in comparing these different bed net technologies and the T-Net impact is actually much greater than shown by these data. The deterrence rate was much lower and the repllency rate much higher for PN2.0 compared to the T-Nets (Table 1, Fig. 3). If the mosquitoes do not enter the but because of deterrence and if they enter the hut but are repelled before receiving a lethal insecticide dose from the bed net, these insects are not being controlled. The model (equation **5**) described in Fig. 4 takes these factors into consideration to determine the “overall relative mosquito control”:

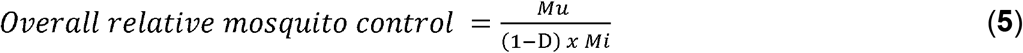

Where:

- *Mu* is the mortality caused by the untreated T-Net (%);
- *Mi* is the mortality caused by the insecticide treated net PN2.0 LLIN (%);
- D is the dettrerent effect 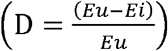
- *Eu* is the mean entry number in untreated T-Net,
- *Ei* is the mean entry number in the insecticide treated PN2.0 LLIN.

From this model, Gen3 was 12.7-fold more efficacious than PN2.0. Gen2 and Gen 1 was 9.1-fold and 7.1-fold, more efficacious (Fig. 4).

## Discussion and conclusion

Malaria is a devastating disease, and vector control is a crucial factor in saving lives. However, the situation has become critical because mosquitoes have become resistant to core intervention tools such as LLINs and IRS. Tackling insecticide resistance by developing new approaches for vector control is imperative to control malaria, and we need to think “out of the box”.

The current model-driven insecticide-free trapping bed-net is a paradigm shift in malaria vector control. Its functionality is based on the attraction and trapping of mosquitoes regardless of their insecticide resistance status (Video 1, suppl. material). This study describes for the first time the efficacy of a trapping bed-net as a malaria vector control strategy, which addresses the problem of insecticide resistance without the need to develop new chemistry. The advantage of the model developed, we can predict the mosquito trapping rate before going into the field for testing; this facilitates additional innovation in the future.

As commonly found in experimental hut trials in which non-insecticide and insecticide-treated products are compared to each-other ^20–21^, we also found higher mosquito entry rates in huts with T-Nets (free of insecticides) than that observed with the PN2.0-LLIN. Similarly, the exit rate was higher for PN2.0-LLIN than that of the T-Nets. This is likely due to the excito-repellency property of insecticides. The excito-repellency effect can be perceived as beneficial to the extent that it keeps some mosquitoes away from the sleeper, and thus provides personal protection to the net user. However, at the community level, the benefit of the excito-repellency effect can be nullified or even contribute to behavioral resistance and increase outdoor malaria transmission as mosquitoes are repelled outdoors and towards unprotected people ^22,23^. Direct observation of field results showed a 2.7 to 4.3-fold increase mortality for the T-Nets comparing to the PN 2.0-LLIN. But, due to the mosquito deterrence as result of excito-repellency, community protection was estimated by our new model to be 13-fold greater for the Gen3 T-Net compared to PN2.0. Thus, mass mosquito trapping as a result of a long-term use of T-Nets should lead to a decline in the vector population to a level where transmission is no longer stable. This population decrease is even more likely in the context of insecticide resistance where mosquito mortality is lower than expected by LLINs.

The T-Net exploits the natural behavior of mosquitoes, which principally interact with the roof of the mosquito net^13,14^. Therefore, the positioning of the traps on the top of the net promotes entrapment of mosquitoes. This is especially true for insecticide-free T-Nets which do not cause repellent effects. Untreated T-Nets can therefore be an asset for vector control as they trap and kill mosquitoes irrespective of their insecticide resistance status. Nevertheless, the addition of a trap compartment to existing LLIN can also enhance the killing performance of the net (Fig. 7, extended data). An insecticide-free tool for malaria vector control is far from being adopted because of regulatory requirements including epidemiological data to prove the public health value, but an interim solution could be a hybrid version of the T-Net consisting of an insecticide-free trap compartment mounted on the currently used LLINs. In this case, community protection would be provided both by the insecticidal effect and the mass trapping of mosquitoes.

Numerous studies have shown that the escalation in resistance to insecticides closely matches the introduction of insecticide-treated nets. To combat resistance to insecticides, the trend today is to use LLINs combined with the synergist piperonyl-butoxide (PBO)^24^. The PBO inhibits enzymatic activity of mosquito insecticide detoxification enzymes, and therefore enhances the insecticidal effect of the treated bed-net. Unfortunately, decreases in performance of these mosquito nets are starting to be observed in some regions of Africa where mosquitoes are resistant^25^. Because the T-Net is insecticide-free, it does not exert any insecticide resistance selection pressure on mosquitoes, and because it is not impacted by insecticide resistance, its exclusive and prolonged use in areas that are endemic for resistance should eventually lead to a decrease in the level of resistance and thus facilitate the reintroduction of insecticides.

Though we have not analyzed the trapped mosquitoes escape rate in the field trial, our lab assays suggest the egression rate was zero. We also have found in laboratory conditions that trapping elicits a mosquito response that rapidly leads to exhaustion and death from dehydration and starvation in one to two days. The dead mosquitoes in the trap compartment are not easily sighted and trapped dead mosquitoes were completely washed out when the T-Nets were hand-washed as per traditional washing methods in Africa. This finding suggests that removing mosquitoes from the T-Net trap compartment is not an issue. They could also be removed by net shaking when the funnels are pulled out. However, we believe that the view of mosquitoes in the trap compartment can be an asset to the extent that it could encourage the use of bed-nets.

Sampling of mosquito populations for transmission studies or evaluation of other vector control tools such as IRS could also be conducted with insecticide-free T-Nets. Various anthropophagous mosquito sampling tools exist but are complex to build up^25–27^. If compared to these methods, the simple construction mode of the T-Net, which uses inexpensive materials, makes it an easy-to-assemble and a less expensive sampling tool. It would be interesting to consider a comparative study of its effectiveness as a sampling method vis-à-vis other methods.

We have not conducted in depth manufacturing cost studies, yet improvements for mass production at low cost are underway. However, we anticipate a satisfactory cost-effectiveness ratio and the ability to stay in the range of 2 to 4 dollars^28^ per T-Net, which is the current range of LLINs costs on the market. Furthermore, because there is no new chemistry to develop, the route to market should be rapid compared to an insecticide treated bed-net. It would also be useful to survey public opinion about the use of an insecticide free bed net versus a LLIN where efficacies for both in a worst-case scenario are equal.

## Materials and Methods

### Tunnel test for assessment of *knitted cone efficiency*

A modified WHO-tunnel^17^ test experiment was conducted to assess the rate at which mosquitoes would move through the knitted cones. The tunnel was modified to include three compartments: the top compartment which operated as a mosquito release chamber, a middle compartment which operated as a trapping container, and a lower compartment that contained the arm of a human subject to attract host-seeking mosquitoes. The top and middle compartments were separated by the knitted cone being tested, and the middle and lower compartment were separated by a piece of bed-net fabric to prevent the human subject from receiving mosquito bites (NCSU IRB approved protocol 16897).

The tunnel was placed vertically to mimic the position of a cone on a T-Net. A human arm was introduced in the bottom compartment. Two replicates of 50 and 106 unfed five to eight-day old host seeking *An. gambiae* Kisumu-strain adult mosquitoes were released into the top compartment. Mosquitoes used were reared at Dearstyne Laboratory according to MR4 rearing protocol^29^. After 2 hours, the mosquitoes that remained in the release chamber were removed using a mouth aspirator and counted. The human arm was removed from the bottom compartment, and trapped mosquitoes were observed for an additional 2 hours. Thereafter, the top compartment was checked for any egression (mosquito passage from the bottom trap compartment to the top compartment). The trapped mosquitoes were then counted.

### Gen3 T-Net proof of concept investigation: laboratory walk-in cage trial

Laboratory walk-in cage trials were conducted in the Dearstyne Entomology Building in the Department of Entomology and Plant Pathology at NCSU in a walk-in sized bioassay cage (1.58 × 2.18 × 2.03 m, depth x width x height) at 27+1°C; 65+4% RH using the Gen3 non-insecticide T-Net. Trapping was conducted with a human subject sleeping under the net (NCSU IRB approved protocol 9067). Five to eight-day old unfed *An. gambiae* Kisumu-strain adult mosquitoes from the same batch above were used. The mosquitoes were released in six replicates of 41 to 106 into the walk-in cage after the human subject had entered the sleeping net compartment. Trapping was conducted for 3 hours during the photophase, since this mosquito strain for years has been fed with lights-on. At the end of the experiment, mosquitoes were vacuum-collected from outside of the trap compartment in the walk-in cage and counted immediately. To assess egression, trapped mosquitoes were not removed from the trap compartment immediately. They were counted immediately after the end of experiment, and after 30 minutes, 1 hour, 2 hours, 6 hours and overnight. They were removed from the trap compartment the following day. The study was conducted on six different days. The mosquitoes were obtained from MR4-BEI Resources, and with current NCSU biosafety approval, a continuous colony was maintained in our laboratory using methods previously described.

### Field study

#### Study site

The study was conducted in experimental huts according to WHO protocol^17^ in the municipality of Tiassalé (5°53’54” N et 4°49’42” W) in November 2018. The site is located in the south of Côte d’Ivoire at about 110 km North of the country’s major city Abidjan. The climate is tropical and characterized by four seasons: a long rainy season (March-July) during which two thirds of the annual rain fall occurs, a short dry season (July-August), a short rainy season (September-November) and a long dry season (December to March). The average annual rainfall is 1,739 mm with an average annual temperature of 26.6 °C. The annual average relative humidity is around 70%. Rice production occurs in the lowlands of Tiassalé which facilitate the proliferation of mosquitoes throughout the year, and malaria is the leading cause of morbidity in the local population. The transmission of the disease is mainly due to *An. coluzzii* (80%) and *An. gambiae* (20%) which have developed multiple resistance to insecticides^19,20.^

#### Experimental station

The experimental field station of Tiassalé is made of 18 standardized experimental huts (Fig. 1a) situated close to the rice plantations. Each hut is 2.5 m long, 1.75 m wide and 2 m high. The walls are made of concrete bricks and plastered with cement, while the floor is made of cement and the roof of corrugated iron sheets. A plastic cover is mounted underneath the roof as a ceiling to facilitate manual collection of mosquitoes. Each hut is built on a platform made of concrete surrounded by a water-filled moat that prevents entry of foraging ants. Entry of mosquitoes is facilitated through four 1 cm wide window slits located on three sides of the hut. The slits are designed in such a way as to prevent mosquitoes from escaping once they are inside the hut. Each hut is equipped with a veranda trap located on the fourth side, made of sheeting and screening mesh to capture mosquitoes that would otherwise escape.

#### Mosquito collection and processing

Four sleepers were trained and paid to sleep under the bed-nets randomly allocated to four huts from 9.00 pm to 5.00 am and to collect mosquitoes the subsequent morning. Sleepers and bed-nets were rotated on a daily basis according to three complete Latin square tables (Fig. 8, extended data) and two additional nights, with the exception of the weekend. Thus, the trial ran for 14 days. Each day after the completion of collections, the huts were cleaned, and the bed-nets rotated appropriately. Prior to the trial, sleepers were all vaccinated against yellow fever and were taken to the hospital to check for malaria parasites. All sleepers gave informed consent prior to enrolment in the study (CSRS IRB protocol #02-2011/MSLS/CNER-P). Within the hut, resting and dead non-trapped mosquitoes were collected from the room and in the veranda trap using 5 ml, 12 mm × 75 mm glass hemolysis tubes. Trapped mosquitoes were removed with mouth aspirators from the T-Net trap compartment by field technicians. All mosquitoes were taken to the field laboratory and identified to genus level and scored as trapped-dead or trapped-alive, not trapped-dead and not trapped-alive (see extended data). Live mosquitoes were placed in cups and given access to sugar solution for 24 h in the insectary at 25-27°C and 70-80% relative humidity (RH) to assess delayed mortality.

#### Study arms

Three different insecticide-free T-Nets (Gen1, Gen2 and Gen3, Fig. 2) were evaluated in comparison to the WHO recommended and widely used Permanet 2.0-LLIN (PN2.0) used as positive control. The PN2.0 was brand new and provided by the National Malaria Control program from Cote d’Ivoire.

#### Data analysis

The Kruskal-Wallis non-parametric test with the Conover-Iman multiple pairwise comparisons and the Bonferroni correction were used to analyze the field data. The precision was fixed at 5%. Analysis was done using the XLSTAT software package version 2019.4.1.^30^ The overall blood-feeding rate was less than 3%; thus, this parameter was not evaluated.

## Supporting information

How does T-Net work

## List of abbreviation

T-Net: Trapping bed-net
Min: minute
WHO: World Health Organization
LLIN: long lasting insecticidal net
PN2.0: Permanet 2.0
Gen: generation
NCSU: North Carolina State University
Std dev.: standard deviation

## Acknowledgments

This work was supported by Innovation Vector Control Consortium (IVCC). CSA and RMR are supported by the NC Ag Res. Service.

## Authors’ contributions

CSM initiate the T-Net conception, contributed to design Gen-1, Gen-2 and Gen-3 T-Nets, carried out laboratory studies, supervised experimental field trials, analysed the data and drafted the manuscript. KL proposed and verified the T-Net model. AW designed and conceived the knitted cones. BKF helped supervising the field study. MMC, MR, CA worked to design the T-Nets, approved the study design, edited and improved the manuscript. All authors read and approved the final manuscript.

## Ethical Approval and Consent to participate

Approval for the research study in experimental huts in Cote d’Ivoire was obtained from the National Ethics Committee (CSRS IRB protocol #02-2011/MSLS/CNER-P). All the sleepers provided written informed consent to participate. Laboratory experiments at NCSU were done in compliance with NCSU IRB approved protocol #16897 and NCSU IRB approved protocol #9067.

## Consent for publication

All the participants consented for publication.

## Availability of data

All data generated or analysed during this study are included in this published article.

## Supplementary information

Other supplementary information is included on extended data file and video file.

## Extended data

### 1. Cone durability assessment

#### Abrasion test method

The Martindale abrasion test method was used to assess the cone textile durability compared to the polyester commonly used bed-net fabric.

Martindale abrasion test refers to the testing of textile products according to Martindale standard system and tests the abrasion resistance of the fabric through the test. Abrasion resistance refers to the resistance of fabric in the process of repeated friction with an abradant.

To run the test, the fabric was loaded onto the lower plates of the Martindale Abrasion machine, and the abradant was then rubbed against the fabric in an oscillating circle. The assay end point was the duration and number of circles until the first appearance of hole. The run speed used was approximately 20 minutes per 1000 circles.

As results, the abrasion duration of cones fabric was 8.67 hours, corresponding to more than 25000 circles, whereas for the bed-net sample, the duration 2.67 hours and less than 10000 circles. Thus, the cone fabric presented a better abrasion resistance than the bed-net fabric. (Figure 5).

**Figure 5:**
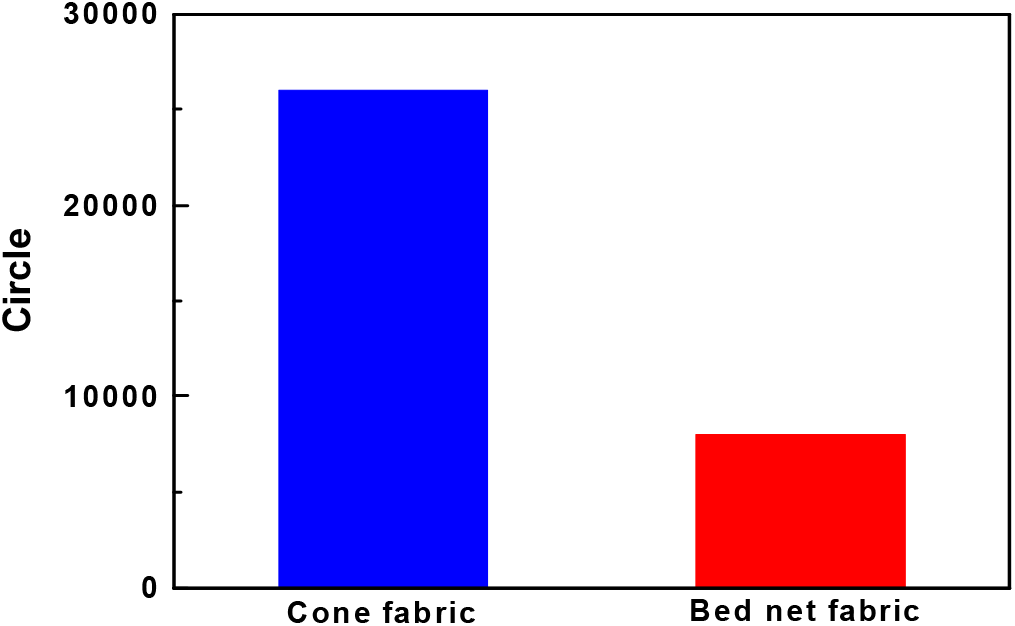
Results obtained after application of Martindale Abrasion Test on a knitted cone fabric compared to conventional polyester 100-dernier bed-net fabric.

#### Burst test

The Burst Test was used to measure the bursting strength of cones and bed-net fabric subjected to an increasing hydrostatic pressure. The pressure is applied to a circular region of the fabric sample via an elastic diaphragm. The fabric sample is firmly held round the edge of this circular region by a pneumatic clamping device. When the pressure is applied, the fabric deforms together with the diaphragm. The bursting strength corresponds to the maximum pressure supported by the fabric before failure.

The test method was ASTM D3786 using the TruBurst machine with the following conditions:

- Conditions: 20°C, 65% RH
- Number of Tests: 5
- Diaphragm: 1.50mm
- Test Area (Dia): 7.3cm^2^ (30.5mm)
- Inflation Rate: 7.67PSI/s
- Correction Rate: 1.45PSI/s
- Burst Detection: Normal
- Clamp Pressure: 44.96PSI

As results, the burst strength, displacement and burst time were higher for the cone fabric than for the bed net textile

**Figure 6:**
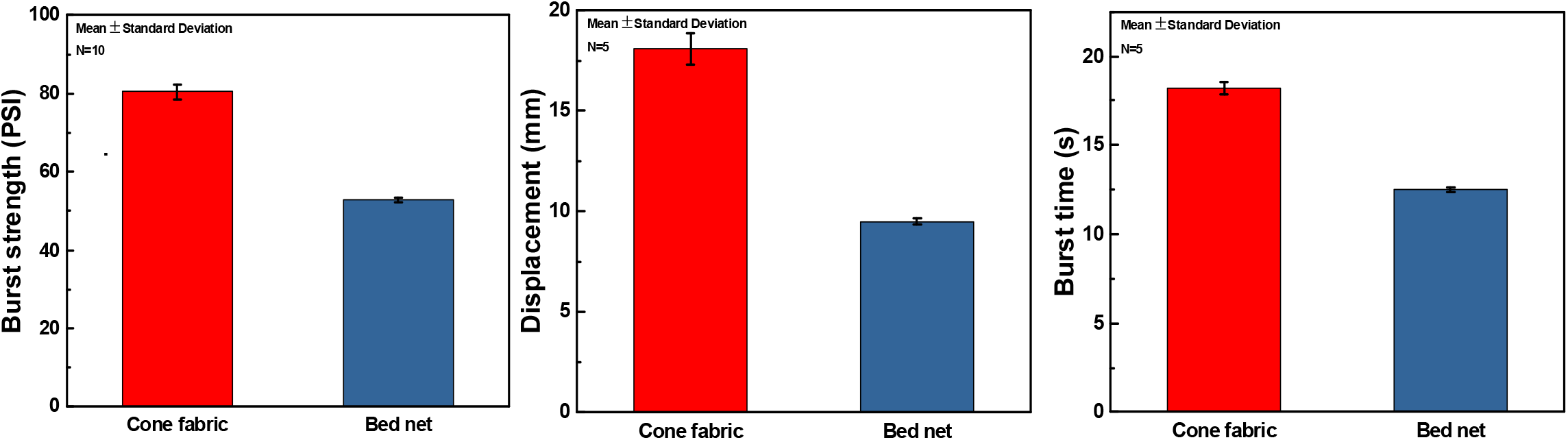
Results obtained after application of Burst Test on a knitted cone fabric compared to conventional polyester 100-dernier bed-net fabric. (a) is the burst strength, (b) is the displacement and (c) is the burst time.

### 2. Long lasting insecticide-treated trapping bed-net efficacy

A preliminary study was carried out in experimental hut to compare the performance of insecticide-free T-Net to insecticide treated T-Net. The methodology and the study sites are the same described in the main body. The only difference is that the T-Net used here were composed of 30 plastic funnels each of the same size (6 cm in the large diameter, 6 cm deep and 2 cm in the small diameter). Here, we compare the trapping potential to the killing activity of insecticide on the same treated T-Net.

As results, we found that about 90% of the mortality registered with the treated T-Net was a result of trapping as most of mosquitoes were found trapped. This results suggest that adding a trap compartment on top of a treated long lasting net can improve its killing potential without impacting the insecticidal activity of the net.

### 3. Latin square table applied for sleepers and bed-net rotations in experimental hut studies

Two different Latin square table designs were used. The first one for the bed-nets rotation and the second one for the sleepers’ rotation (Figure 7). Both were used simultaneously to give a chance to sleeper to occupy each hut and utilize each bed-net.

**Figure 7.**
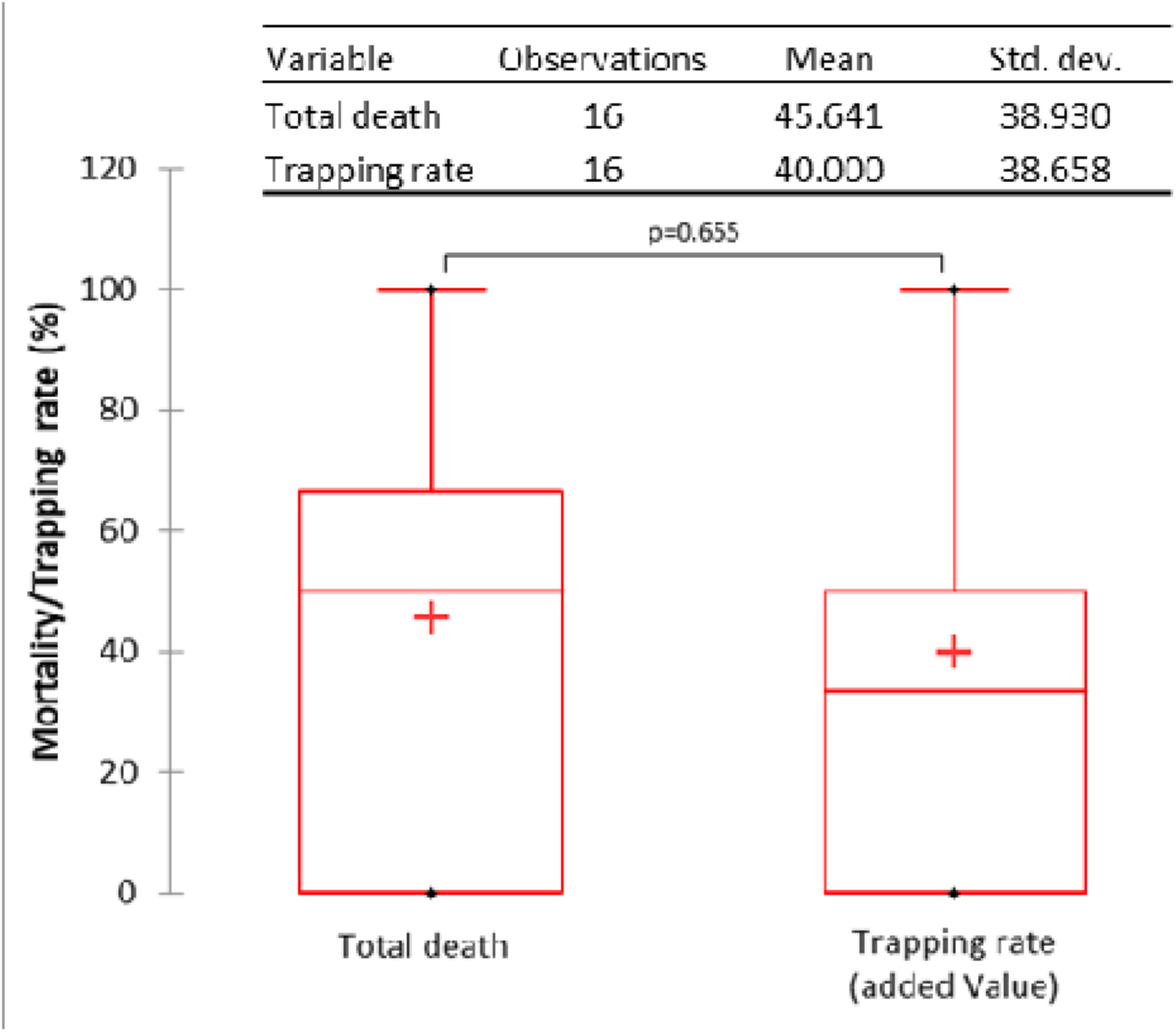
Overall mortality versus the trapping rate with the insecticide-treated T-Net. The horizontal bar indicates the P-value obtained with the Mann-Whitney statistical test. The table above shows the mean values and the standard deviations after 16 observations.

**Figure 8.**
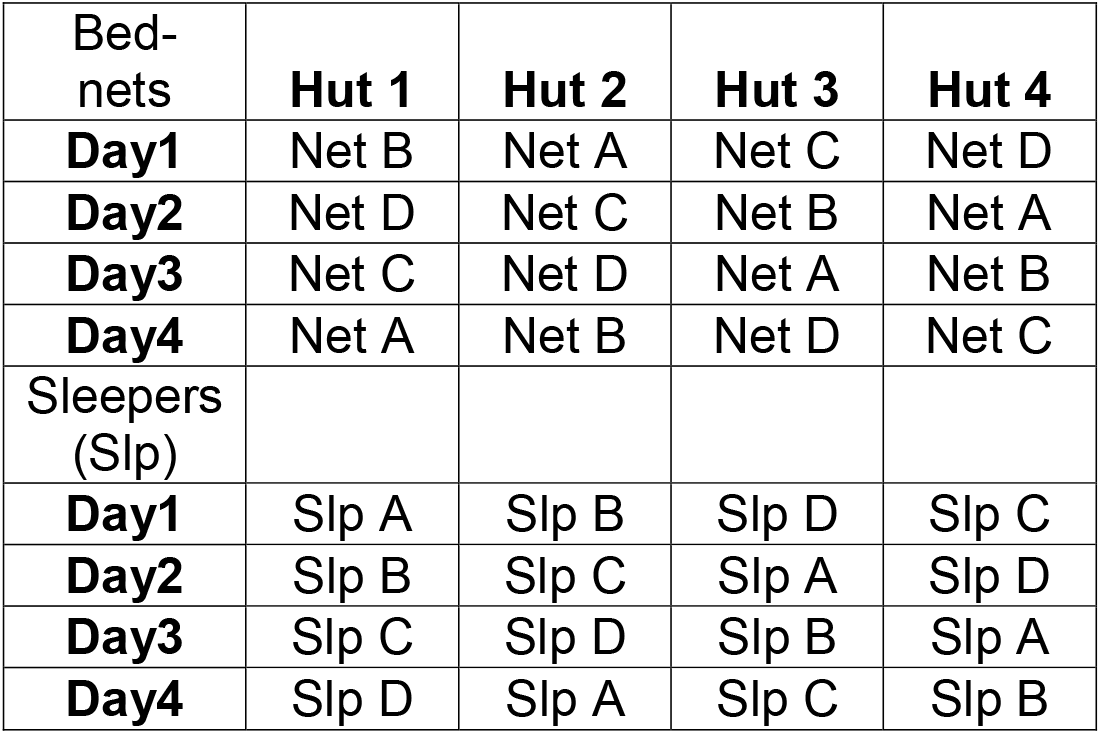
Ballot box draws were used to attribute a letter to each bed-net as well as to each sleeper. Bed-nets and sleepers were then rotated according the above order generated by a Latin square table generator. Every four days, new other ballot box draws were carried out to redistribute letters to the bed-nets and to the sleepers.

## References

1. World malaria report 2016. (2016).

2. Global plan for insecticide resistance management in malaria vectors. (World Health Organization, 2012).

3. Bhatt, S. et al. The effect of malaria control on Plasmodium falciparum in Africa between 2000 and 2015. Nature 526, 207–211 (2015).

4. Ranson, H. & Lissenden, N. Insecticide Resistance in African Anopheles Mosquitoes: A Worsening Situation that Needs Urgent Action to Maintain Malaria Control. Trends in Parasitology 32, 187–196 (2016).

5. Müller, P. et al. Pyrethroid tolerance is associated with elevated expression of antioxidants and agricultural practice in Anopheles arabiensis sampled from an area of cotton fields in Northern Cameroon: INSECTICIDE TOLERANCE IN ANOPHELES ARABIENSIS. Molecular Ecology 17, 1145–1155 (2007).

6. Edi, C. V. A., Koudou, B. G., Jones, C. M., Weetman, D. & Ranson, H. Multiple-Insecticide Resistance in *Anopheles gambiae* Mosquitoes, Southern Côte d’Ivoire. Emerg. Infect. Dis. 18, 1508–1511 (2012)

7. Strode, C., Donegan, S., Garner, P., Enayati, A. A. & Hemingway, J. The Impact of Pyrethroid Resistance on the Efficacy of Insecticide-Treated Bed Nets against African Anopheline Mosquitoes: Systematic Review and Meta-Analysis. PLoS Med 11, e1001619 (2014).

8. Hemingway, J. The role of vector control in stopping the transmission of malaria: threats and opportunities. Phil. Trans. R. Soc. B 369, 20130431 (2014).

9. Ranson, H. et al. Pyrethroid resistance in African anopheline mosquitoes: what are the implications for malaria control? Trends in Parasitology 27, 91–98 (2011).

10. https://www.ivcc.com/research-development/insecticide-discovery-and-development/

11. Mouhamadou, C. S. et al. Evidence of insecticide resistance selection in wild Anopheles coluzzii mosquitoes due to agricultural pesticide use. Infect Dis Poverty 8, 64 (2019).

12. Van Loon, J. J. A. et al. Mosquito Attraction: Crucial Role of Carbon Dioxide in Formulation of a Five-Component Blend of Human-Derived Volatiles. J Chem Ecol 41, 567–573 (2015).

13. Parker, J. E. A. et al. Infrared video tracking of Anopheles gambiae at insecticide-treated bed nets reveals rapid decisive impact after brief localised net contact. Sci Rep 5, 13392 (2015).

14. Sutcliffe, J., Ji, X. & Yin, S. How many holes is too many? A prototype tool for estimating mosquito entry risk into damaged bed nets. Malar J 16, 304 (2017).

15. Hernandez, H. Standard Maxwell-Boltzmann distribution: Definition and Properties. (2017) doi:10.13140/RG.2.2.29888.74244.

16. Darriet, F. & N’Guessan, R. Un outil expérimental indispensable à l’évaluation des insecticides◻: les cases-pièges. Bull Soc Pathol Exot 6 (2002).

17. WHO_CDS_WHOPES_GCDPP_2005.11.pdf.

18. Chouaïbou, M. S. et al. Influence of the agrochemicals used for rice and vegetable cultivation on insecticide resistance in malaria vectors in southern Côte d’Ivoire. Malar J 15, 426 (2016). Fodjo, B. K. et al. Insecticides Resistance Status of *An. gambiae* in Areas of Varying Agrochemical Use in Côte D’Ivoire. BioMed Research International 2018, 1–9 (2018).

19. Fodjo, B. K. et al. Insecticides Resistance Status of *An. gambiae* in Areas of Varying Agrochemical Use in Côte D’Ivoire. BioMed Research International 2018, 1–9 (2018).

20. Menze, B. D. et al. An Experimental Hut Evaluation of PBO-Based and Pyrethroid-Only Nets against the Malaria Vector Anopheles funestus Reveals a Loss of Bed Nets Efficacy Associated with GSTe2 Metabolic Resistance. Genes 11, 143 (2020).

21. Ngufor, C. et al. Combining Organophosphate Treated Wall Linings and Long-lasting Insecticidal Nets for Improved Control of Pyrethroid Resistant Anopheles gambiae. PLoS ONE 9, e83897 (2014).

22. Russell, T. L., Beebe, N. W., Cooper, R. D., Lobo, N. F. & Burkot, T. R. Successful malaria elimination strategies require interventions that target changing vector behaviours. Malaria Journal 12, 56 (2013).

23. Sougoufara, Seynabou, et al. “Challenges for malaria vector control in sub-Saharan Africa: Resistance and behavioral adaptations in anopheles populations.” Journal of Vector Borne Diseases, vol. 54, no. 1, 2017, p. 4. Gale Academic Onefile, Accessed 30 Jan. 2020.

24. Toé, K. H. et al. Increased Pyrethroid Resistance in Malaria Vectors and Decreased Bed Net Effectiveness, Burkina Faso. Emerg. Infect. Dis. 20, (2014).

25. Govella, N. J. et al. A new tent trap for sampling exophagic and endophagic members of the Anopheles gambiae complex. Malar J 8, 157 (2009).

26. LaBrecque, B. et al. Sampling Host-Seeking Anthropophilic Mosquito Vectors in West Africa: Comparisons of an Active Human-Baited Tent-Trap Against Gold Standard Methods. The American Journal of Tropical Medicine and Hygiene 92, 415–421 (2015).

27. Tangena, J.-A. A., Thammavong, P., Hiscox, A., Lindsay, S. W. & Brey, P. T. The Human-Baited Double Net Trap: An Alternative to Human Landing Catches for Collecting Outdoor Biting Mosquitoes in Lao PDR. PLoS ONE 10, e0138735 (2015).

28. https://www.unicef.org/supply/files/Long-Lasting_Insecticidal_Nets_price_data_January_2018.pdf

29. Benedict, M. Q. Methods in Anopheles research. Malaria Research and Reference Reagent Resource Center (MR4) (2007).

30. Addinsoft (2020). XLSTAT 2019.4.1 statistical and data analysis solution. New York, USA. https://www.xlstat.com

